# A stochastic model of gene expression with polymerase recruitment and pause release

**DOI:** 10.1101/717199

**Authors:** Z. Cao, T. Filatova, D. A. Oyarzún, R. Grima

## Abstract

Transcriptional bursting is a major source of noise in gene expression. The telegraph model of gene expression, whereby transcription switches between “on” and “off” states, is the dominant model for bursting. Recently it was shown that the telegraph model cannot explain a number of experimental observations from perturbation data. Here we study an alternative model that is consistent with the data and which explicitly describes RNA polymerase recruitment and polymerase pause release, two steps necessary for mRNA production. We derive the exact steady-state distribution of mRNA numbers and an approximate steady-state distribution of protein numbers which are given by generalized hypergeometric functions. The theory is used to calculate the relative sensitivity of the coefficient of variation of mRNA fluctuations for thousands of genes in mouse fibroblasts. This indicates that the size of fluctuations is mostly sensitive to the rate of burst initiation and the mRNA degradation rate. Furthermore we show that (i) the time-dependent distribution of mRNA numbers is accurately approximated by a modified telegraph model with a Michaelis-Menten like dependence of the effective transcription rate on RNA polymerase abundance. (ii) the model predicts that if the polymerase recruitment rate is comparable or less than the pause release rate, then upon gene replication the mean number of RNA per cell remains approximately constant. This gene dosage compensation property has been experimentally observed and cannot be explained by the telegraph model with constant rates.

**Statement of Significance:** The random nature of gene expression is well established experimentally. Mathematical modelling provides a means of understanding the factors leading to the observed stochasticity. There is evidence that the classical two-state model of stochastic mRNA dynamics (the telegraph model) cannot describe perturbation experiments and a new model that includes polymerase dynamics has been proposed. In this paper, we present the first detailed study of this model, deriving an exact solution for the mRNA distribution in steady-state conditions, an approximate time-dependent solution and showing the model can explain gene dosage compensation. As well, we use the theory together with transcriptomic data, to deduce which parameters when perturbed lead to a maximal change in the size of mRNA fluctuations.

## 1 Introduction

There is widespread evidence that mammalian genes are expressed in bursts: infrequent periods of transcriptional activity that produce a large number of messenger RNA (mRNA) transcripts within a short period of time [1–3]. This is in contrast to constitutive expression where mRNAs are produced in random, uncorrelated events, with a time-independent probability [4]. The size and frequency of transcriptional bursts affect the magnitude of temporal fluctuations in mRNA and protein content of a cell, and thus constitute an important source of intracellular noise [5].

A large number of studies have sought to elucidate the mechanisms leading to bursting and by constructing simple stochastic models that can explain the data. The simplest of these models is the telegraph model whereby (i) a gene is in two states, an ON state where mRNA is expressed and an OFF state where there is no expression. (ii) mRNA degrades in the cytoplasm. These first-order reactions are effective since each encapsulates the effect of a large number of underlying biochemical reactions. The chemical master equation of this model has been solved exactly to obtain the probability distribution of mRNA numbers as a function of time [6]. For parameter conditions consistent with bursty expression, the steady-state distribution is well approximated by a negative binomial that fits some of the experimental data [7].

Recent studies have extended the telegraph model in various directions (see [8] for a recent review). Mammalian cells have been shown to display complex promoter dynamics during the switch from transcriptionally inactive to active states. Such dynamics cannot be described by a single reaction step whose time is exponentially distributed [2], as assumed by the telegraph model. In [9] this complexity is accounted for by deriving analytical expressions linking the Fano factor of mRNA distributions to the general waiting-time distribution of the time to switch from inactive to active states. In contrast, other works [10–12] have sought to describe promoter dynamics with transitions between a number of discrete promoter states, only some of which are active; in special cases of such models, the steady-state distribution of mRNA fluctuations can be derived analytically. Moreover, dynamic regulation of eve stripe 2 expression in living Drosophila [13] suggests the occurrence of multiple rates of Pol II loading, which argues in favour of the multistate model rather than the simpler telegraph model. Another study, based on live cell imaging of the amoeba Dictyostelium, postulates a continuum of transcriptional states [14] rather than discrete states. All these models share a common property with the telegraph model, namely that when a transcript is produced, the gene state is unchanged.

Bartman et al. [15] recently argued that it is unclear how polymerase recruitment and pause release, two well-known steps in mRNA production, map onto the active and inactive states assumed by the telegraph model. This argument also applies to the various multistate variants of the telegraph model. In particular, in these models one cannot tell whether the initiation of a burst permits polymerase recruitment to occur or whether it permits release from the paused state. In [15], the telegraph model and several possible models of transcription were considered that incorporated bursting (burst initiation and termination steps) together with polymerase recruitment and pause release steps. Using stochastic simulations in conjunction with RNA FISH and Pol II ChIP-seq measurements, they showed that the only model compatible with the data is one in which (i) polymerase recruitment follows after burst initiation and (ii) only one polymerase is permitted to bind each promoter-proximal region at a time, and this bound polymerase has to undergo pause release before a second polymerase can be recruited to a gene copy (in line with the findings in [16, 17]). We emphasize that while this model has three effective gene states, it is not a special case of the multistate gene models studied in [10–12]. These models assume that the gene state does not change upon production of mRNA because they model the production of a mature transcript without detailed modelling of the steps between transcriptional initiation and termination. However the model expounded in [15] models transcription at a finer level of detail which requires that the production of nascent mRNA results in a change of gene state, a property that is crucial to capture property (ii) above. Note the number of nascent mRNA molecules, irrespective of their length, is equal to the number of polymerases currently transcribing the gene [18]. An interesting recent review discussing the assumptions behind common gene expression models including those with polymerase dynamics can be found in [19].

In this article, we present the first detailed study of the model proposed by Bartman et al [15]. The article is organized as follows. In Section 2.1 we introduce the chemical master equation formulation of the model, in Section 2.2 we obtain an an exact steady-state solution of this model and in Section 2.3 we use the theoretical results and transcriptomic data to investigate the sensitivity of the size of mRNA fluctuations to the five parameters. In Section 2.4 we show that by mapping the model onto an effective telegraph model, we can obtain an approximate time-dependent solution. In Section 2.5 we show that while our model has three effective promoters states, it is not the same as the refractory model of gene expression devised by Naef and co-workers [2]. In Section 2.6 we show that the protein number distribution can also be obtained in the limit of fast mRNA decay and that this is generally different than that obtained using the conventional three-stage model of gene expression [20]. We finish with a discussion of the biological implications of our results in Section 3.

## 2 Results

### 2.1 Model

We consider a stochastic transcriptional bursting model (recently introduced in [15] and henceforth referred to as the multi-scale model; see Fig. 1A), whereby a gene fluctuates between three states: two permissive states (*D*_10_ and *D*_11_) and a non-permissive state (*D*_0_).

**Figure 1:**
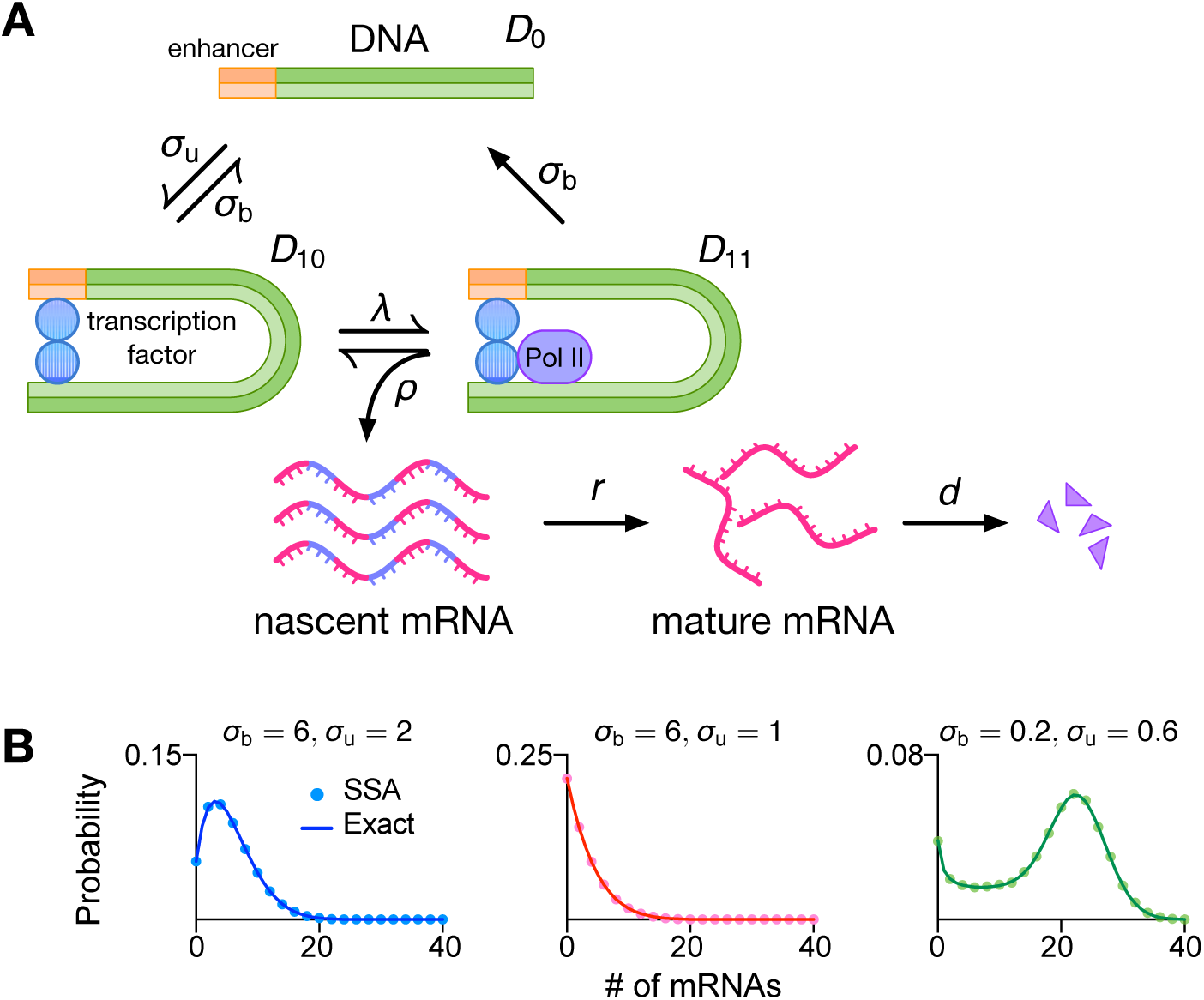
(A) Schematic of the stochastic multi-scale transcriptional bursting model. (B) Analytical distribution for mature mRNA numbers (under the assumption of short-lived nascent mRNA) is given by Eq. (6) and agrees with stochastic simulations using the SSA. The kinetic parameters are *ρ* = 60, *λ* = 40, *d* = 1; other parameters are indicated in each panel.

The transition from *D*_0_ to *D*_10_ (burst initiation) is mediated by transcription factor binding with rate constant *σ*_u_ which is reversible with rate constant *σ*_b_ (this transition may alternatively represent other processes such as nucleosome remodeling). Subsequently the binding of RNA polymerase II (Pol II) to *D*_10_ with rate constant *λ* (which is proportional to Pol II abundance) leads to *D*_11_. This represents a state in which Pol II is paused, and models the experimental observation that Pol II pauses downstream of the transcription initiation site preceding productive elongation [17]. The polymerase is released from this state with rate constant *ρ* leading to two simultaneous processes: (i) since now the polymerase can actively transcribe RNA, it implies the production of nascent mRNA (denoted as *N*) with rate *ρ*; (ii) the gene state changes from *D*_11_ to *D*_10_. This step models the experimental observation that unless the polymerase is unpaused, there is no binding of new Pol II [16, 17]. In the paused state *D*_11_, both the polymerase and the transcription factor can unbind from the gene and lead to the non-permissive state *D*_0_ (burst termination). Both reversible switches operate at different timescales (hours versus minutes) with max{*σ*_b_, *σ*_u_} ≪ min{*ρ, λ*}, leading to multi-scale transcriptional bursting [15, 21]. After termination, the nascent mRNA becomes a mature mRNA (denoted by *M*); this occurs with rate *r*. Subsequently the mature mRNA decays with rate constant *d*. Note that we assume all reactions to be first-order, characterized by exponentially distributed waiting times between successive reactions.

In what follows, for simplicity, we assume that the lifetime of nascent mRNA is very short, i.e. *r* is large, such that the reaction *D*_11_ → *D*_10_ + *N, N* → *M* can be approximated by the single reaction step *D*_11_ → *D*_10_ + *M*. In the next section, we derive the steady-state distribution of mature mRNA (simply called mRNA henceforth).

### 2.2 Exact solution

Let *P*_*θ*_(*n, t*) (*θ* = 0, 10, 11) denote the probability of a cell being in state *D*_*θ*_ with *n* mRNAs at time *t* (arguments *n* and *t* are hereafter omitted for brevity). The dynamics of probability *P*_*θ*_ are described by the set of coupled master equations

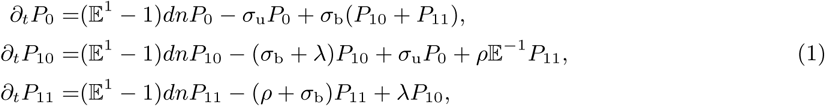

where the step operator 𝔼^*i*^ acts on a general function *g*(*n*) as 𝔼^*i*^*g*(*n*) = *g*(*n* + *i*) [22]. To solve Eq. (1), we use the generating function method and define *G*_*θ*_(*z*) = Σ_*n*_ *z*^*n*^*P*_*θ*_(*n*) for *θ* = 0, 10, 11 so that Eq. (1) can be recast as a set of coupled partial differential equations

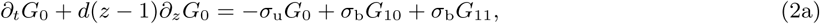

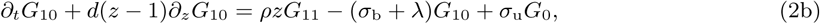

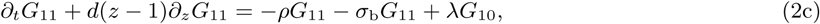

wherein the variable *z* is dropped for brevity. By setting *z* = 1 and the time derivatives to zero (considering steady-state conditions), we can deduce that the probability of being in the non-permissive state *D*_0_ is *G*_0_(1) = *σ*_*b*_*/*(*σ*_*u*_ + *σ*_*b*_) and the probability of being in one of the two permissive states *D*_10_ or *D*_11_ is *G*_10_(1) + *G*_11_(1) = *σ*_*u*_*/*(*σ*_*u*_ + *σ*_*b*_).

In order to solve Eq. (2) for *G*_0_(*z*), *G*_10_(*z*), *G*_11_(*z*) in steady-state conditions, we set *∂*_*t*_*G*_*θ*_ = 0, solve *G*_10_ from Eq. (2c) as a function of *G*_11_, and combine the yielded result to solve *G*_0_ from Eq. (2b) as a function of *G*_11_ so that Eq. (2a) consequently becomes a differential equation with *G*_11_ being the only variable

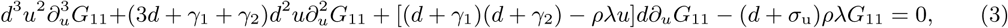

with *u* = *z* − 1, *γ*_1_ = *σ*_b_ + *σ*_u_ and *γ*_2_ = *ρ* + *λ* + *σ*_b_. By defining a new variable *x* = *ρλu/d*^2^, Eq. (3) can be further simplified to

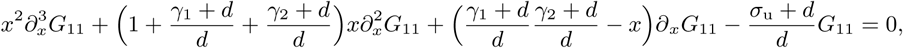

which is in the canonical form of the differential equation for the generalized hypergeometric function

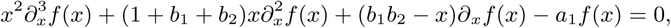

admitting the solution *f* (*x*) = *C*_1_*F*_2_(*a*_1_; *b*_1_, *b*_2_; *x*) with *C* being an integration constant. Hence, the solution for *G*_11_ is in terms of the generalized hypergeometric function

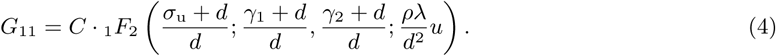

On the other hand, summing Eqs. (2a)-(2c) and denoting *G* = Σ_*θ*_ *G*_*θ*_, one can get *∂*_*u*_*G* = *ρG*_11_*/d*, which together with Eq. (4) leads to

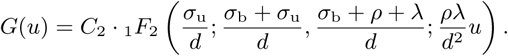

Note that in the last step we made use of the general relation 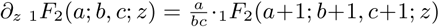. The integration constant *C*_2_ is found to be 1 by using the normalization condition *G*(0) = 1. Hence, the exact solution for the generating function is

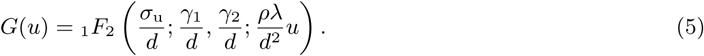

Hence it follows that the marginal probability of finding *n* mRNAs in a cell is

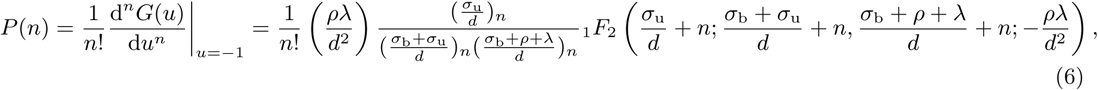

where (·)_*n*_ is the Pochhammer symbol. In Fig. 1B we show that distributions obtained from Eq. (6) as well as the corresponding modality (a phenotypic signature [23]) are indistinguishable from distributions produced using the stochastic simulation algorithm (SSA) [24]. Note that here we have solved for the mature mRNA distribution, under the assumption that nascent mRNA is short lived. In cases where this assumption is not physiologically meaningful and one is interested in the nascent mRNA distribution, then the latter is given by Eq. 6 with *d* replaced by *r* (the rate at which nascent mRNA changes to mature mRNA due to the termination of transcription).

#### 2.2.1 Special case of bursty transcription

It can be further shown by perturbation theory in Appendix A that when *ρ, λ* and *σ*_b_ are much greater than the rest of the parameters, the exact solution Eq. (6) reduces to the negative binomial distribution 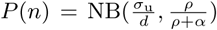 with *α* = *σ*_b_*γ*_2_*/λ*. Note the constraint on the parameters leads to time-series with large and short-lived bursts of transcription (since *ρ, λ*, and *σ*_*b*_ are large), separated by long silent intervals (since *σ*_*u*_ is small). Such bursty trancription is common in mammalian cells [3].

#### 2.2.2 Relationship to the telegraph model

It can also be shown that that in the limit of large *ρ*, the exact solution Eq. (6) reduces to the confluent hypergeometric solution of the telegraph model (see Appendix B). This is equivalent to the steady-state solution of the two state system 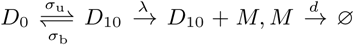. The reduction to a two state model results from genes spending a short time in state *D*_11_ due to the large value of *ρ*. The production of an mRNA molecule involves the slow reaction step from *D*_10_ to *D*_11_ with rate *λ* followed by a very fast reverse step with rate *ρ*. Hence the rate of mRNA production is determined by the reaction rate of the slowest reaction, i.e. it is equal to *λ*. By similar reasoning, we can deduce that in the limit of large *λ*, the gene spends short time in the state *D*_10_ and the multi-scale model reduces to the two state telegraph model with a rate of mRNA production equal to *ρ*.

### 2.3 Sensitivity analysis

The exact solution in Eq. (5) allows us to examine the stochastic properties of the multi-scale model over large swathes of parameter space. We investigate the relative sensitivity of the coefficient of variation of mRNA fluctuations, 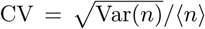, which is typically employed as a measure of the magnitude of transcriptional noise. To this end, we calculate the first two central moments, (⟨*n*⟩ and Var(*n*)), from Eq. (5) using ⟨*n*⟩ = *∂*_*u*_*G*|_*u*=0_ and Var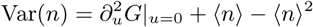. The mean and CV are then given by

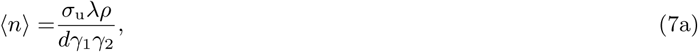

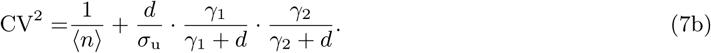

Note that since the parameters *ρ* and *λ* appear symmetrically in Eq. (7), for simplicity we enforce the constraint *ρ* = *λ* (we will relax this constraint later). Hence, the relative sensitivity of the quantity 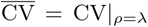, which can serve as a gauge of transcriptional noise, is insightful to study and defined as 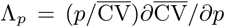 for a model parameter *p*, meaning that 1% change in *p* leads to a Λ_*p*_% change in 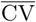. The parameter values for the sensitivity analysis were sampled from experimental distributions recently inferred for 3575 genes of CAST allele in mouse fibroblasts [3], using the telegraph model. To obtain values for *ρ* and *λ*, we equate the mean of the telegraph model (with ON switching rate *σ*_b_, OFF switching rate *σ*_u_, transcription rate *ρ*_u_ and degradation rate *d*) ⟨*n*⟩_tel_ = *σ*_u_*ρ*_u_*/γ*_1_*d* with the mean of the multi-scale model (Eq. (7a)) under the constraint *ρ* = *λ*, giving

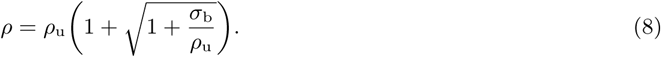

Distributions for each parameter in the dataset are presented in Fig. 2A and the box plots in Fig. 2B show the relative sensitivity for each parameter. The parameters in order of most sensitive first are *σ*_u_, *d, σ*_b_ and *ρ* = *λ*. This order is the same as obtained by ranking parameters according to the inverse of their mean experimental values (the mean of the distributions in Fig. 2A) implying that changes to the CV are most easily accomplished by perturbations to the slowest reactions. Given the vectors Λ_*p*_1 and Λ_*p*_2 for any pair *p*_1_ ≠ *p*_2_ and *p*_1_, *p*_2_ in the set {*ρ, λ, σ*_b_, *σ*_u_, *d*} where each entry is a different gene, in Fig. 2C we calculate the Pearson correlation coefficient between the vectors and the corresponding joint distributions. This shows that (*σ*_u_, *σ*_b_) is the least dependent pairing and hence they constitute a quasi-orthogonal decomposition of the sensitivity. In other words, a change in the CV due to a change in *σ*_u_ is practically uncorrelated with a change in the CV due to a change in *σ*_b_, and hence these two parameters can be seen as independent “control knobs” to change the CV; this is of interest in synthetic biology, where an engineering design approach is taken to modify a biological system for improved functionality [25, 26]. The same set of parameters ranked by sensitivity are obtained, if instead of setting *λ* = *ρ*, we consider *ρ* ≫ *λ*, or *λ* ≫ *ρ*, and hence it appears that our results in this section are robust and invariant with respect to the ratio *λ/ρ*.

**Figure 2:**
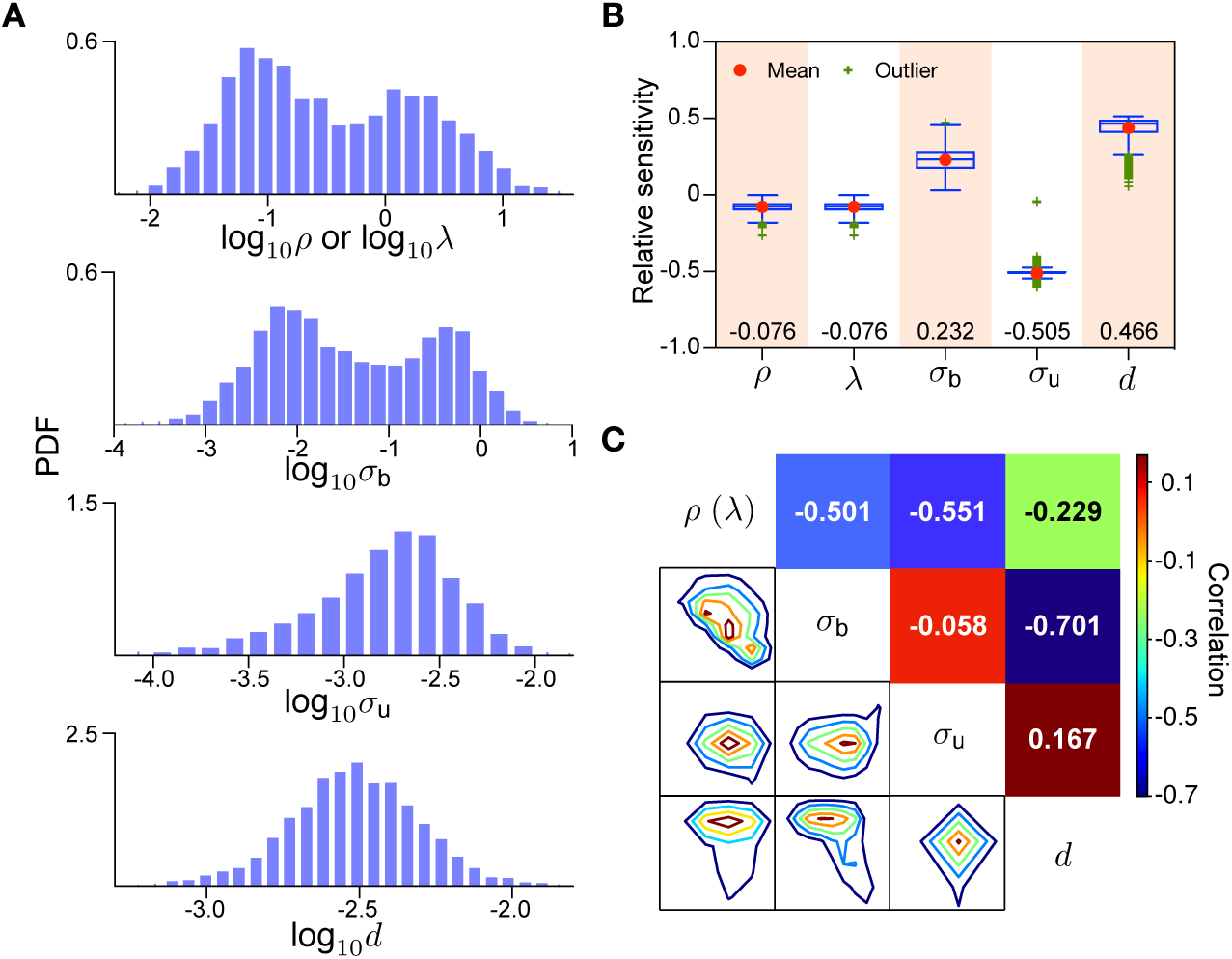
Relative sensitivity analysis of the coefficient variation 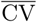 of mRNA noise over 5 kinetic parameters for 3575 genes of CAST allele data for mouse fibroblasts.(A) Distributions of the kinetic parameters in the dataset (obtained from [3]); values of *ρ* or *λ* are calculated using Eq. (8). (B) Box plots indicate the median (values shown at bottom), the 25%, 75% quantiles, and mean and outliers of relative sensitivity. (C) Joint distributions and Pearson correlation between the relative sensitivity vectors for each pair of parameters suggest that (*σ*_b_, *σ*_u_) and (*σ*_u_, *d*) are the least dependent pairs.

### 2.4 Effective telegraph model

Earlier we showed that in the limit of large *ρ* or large *λ*, the solution of the multi-scale model tends to the solution of the telegraph model. Next we use the first passage time method to reduce the multi-scale model into an effective telegraph model, without making the aforementioned assumptions. To this end, we consider the transcription motif of the multi-scale model, 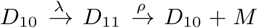, whose corresponding master equations for producing newborn mRNA starting from state *D*_10_ are

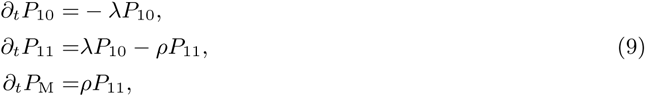

where *P*_10_, *P*_11_ and *P*_M_ represent the probability of staying in states *D*_10_, *D*_11_ or producing a new mRNA respectively. We remark that the reaction *D*_11_ → *D*_0_ is absent from the motif due to its relatively small reaction rate *σ*_b_ compared to *ρ* and *λ*. The initial conditions for Eq. (9) are *P*_10_|_*t*=0_ = 1, *P*_11_|_*t*=0_ = *P*_M_|_*t*=0_ = 0. Solving for *P*_M_ in Eq. (9), we can calculate the mean first passage time for mRNA production

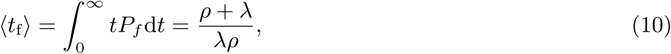

where *P*_*f*_ = *∂*_*t*_*P*_M_ is the first-passage time distribution [27]. Since the effective transcription rate is the inverse of the mean first passage time, it immediately follows that the effective telegraph model is

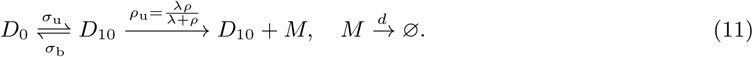

Alternatively, one can obtain this result by equating the means of our model Eq. (7a) and of the telegraph model ⟨*n*⟩_tel_ = *ρ*_u_*σ*_u_*/γ*_1_*d* and solving for the effective production rate *ρ*_u_, giving 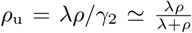 since typically *ρ, λ* ≫ *σ*_b_.

In Fig. 3, we show the high accuracy of the effective telegraph model approximation from Eq. (11). In particular, Fig. 3A shows a heatmap of the distance between the distributions of mRNA numbers predicted by the effective telegraph model and the multi-scale model. As a distance measure, we use the Hellinger distance (HD), a Euclidean distance based metric normalized to the interval between 0 and 1. The effective telegraph model is naturally a more accurate description to the multi-scale model when there is one rate limiting step (large difference between *ρ* and *λ*) rather than when there are two rate limiting steps (*ρ* = *λ*).

**Figure 3:**
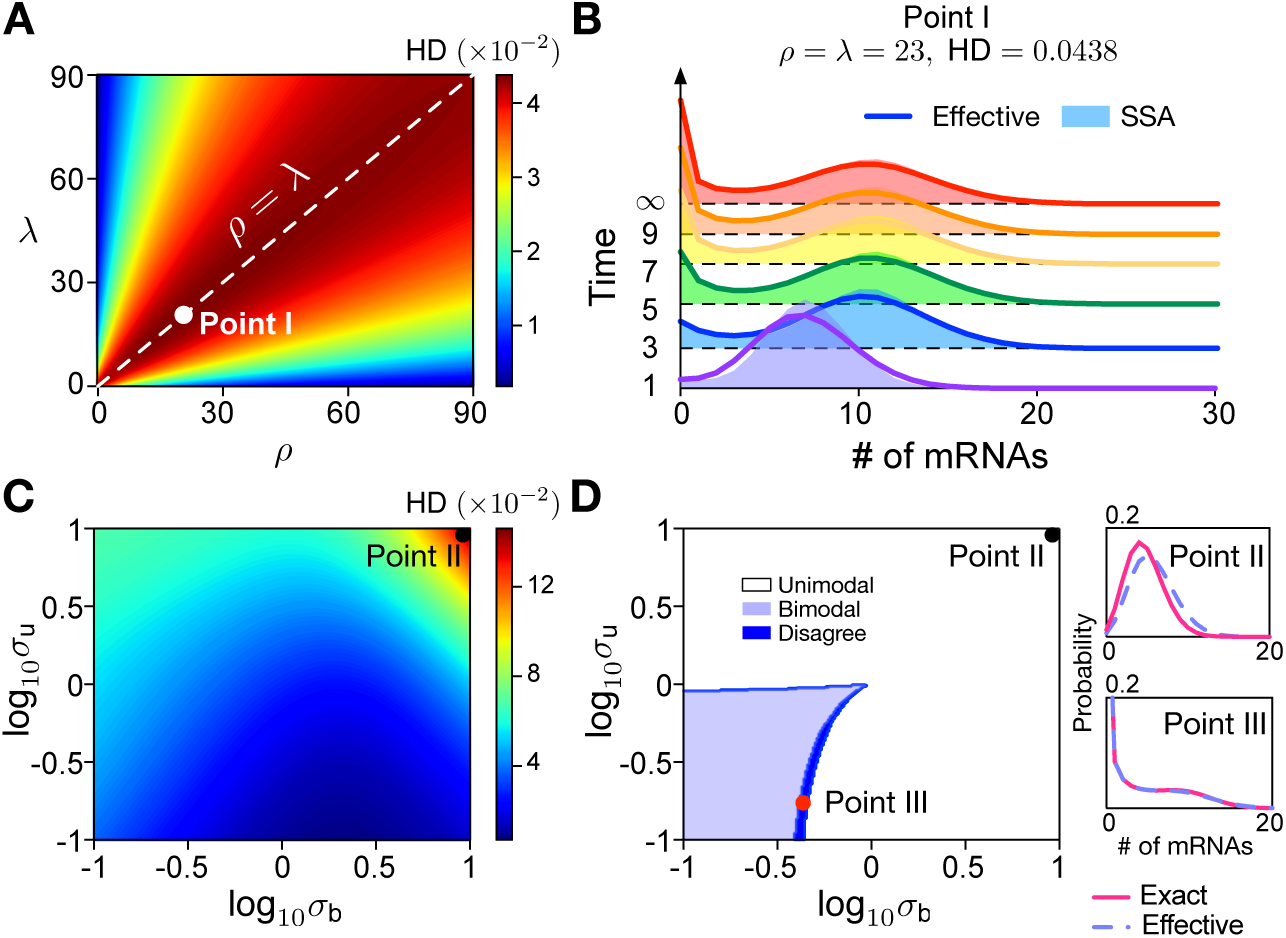
An effective telegraph model (given by reaction scheme (11)) approximates the distribution of mRNA numbers of the multi-scale model. (A) Hellinger Distance (HD) between steady-state distributions of mRNA numbers for the effective telegraph model and the multi-scale model as a function of *ρ* and *λ* with *σ*_u_ = 0.2, *σ*_b_ = 0.1 and *d* = 1. The discrepancy between the two distributions grows as *ρ* and *λ* approach the line *ρ* = *λ*. (B) Shows the time-dependent distributions for Point I in (A) (the point with the largest HD) predicted by the effective model compared to those computed by the SSA for the multi-scale model. (C) Heatmap of HD between both distributions as a function of *σ*_b_ and *σ*_u_ with *ρ* = *λ* = 23 and *d* = 1. (D) Stochastic bifurcation diagram for the number of modes of the steady-state distributions predicted by the two models. The small dark blue region is where modality of both models disagree. Insets show distributions corresponding to the points marked in (C,D).

Since the time-dependent distribution of the telegraph model is known in closed-form [6, 28], it follows that by the effective model in Eq. (11) we have an approximation for the time-dependent distribution of the multi-scale model too. The accuracy of this approximation is shown in Fig. 3B where it is compared to the time-dependent distributions computed using the SSA for the multi-scale model. The parameters here correspond to those of Point I in Fig. 3A (the largest HD). Differences between the distributions of the two models are negligible except near time *t* = 0. We further investigate how burst initiation and termination rates (*σ*_u_, *σ*_b_) affect the approximation error with a heatmap of HD as a function of *σ*_u_ and *σ*_b_ (Fig. 3C), and a stochastic bifurcation diagram for the number of modes of the effective telegraph and multi-scale model distributions (Fig. 3D) at steady state. The point of maximum HD in Fig. 3C (Point II) displays distributions that are not that different from each other – see upper right inset of Fig. 3D. The two models display the same number of modes in all regions of parameter space except for a narrow region where modality detection is challenging because the distributions have a broad plateau – see lower right inset of Fig. 3D (Point III). This again confirms the high accuracy of the effective telegraph model approximation. The biological implications of the Michaelis-Menten dependence of the transcription rate *ρ*_u_ in Eq. (11) on *λ* and *ρ* is discussed in the Conclusion; in particular there we argue how this special feature of our model can explain gene dosage compensation observed in experiments.

### 2.5 Connection to the refractory model

Besides the telegraph model, another prevalent stochastic transcriptional model is the refractory model [2] (a three-state model, see Fig. 4A left), wherein the burst initiation requires two steps. This model was devised to explain the experimental observation that the distribution of “off” intervals is not exponential but rather has a peak at a non-zero value. To understand the connection between our model and the refractory model, we first exactly solve the refractory model for the steady-state distribution of mRNA numbers.

**Figure 4:**
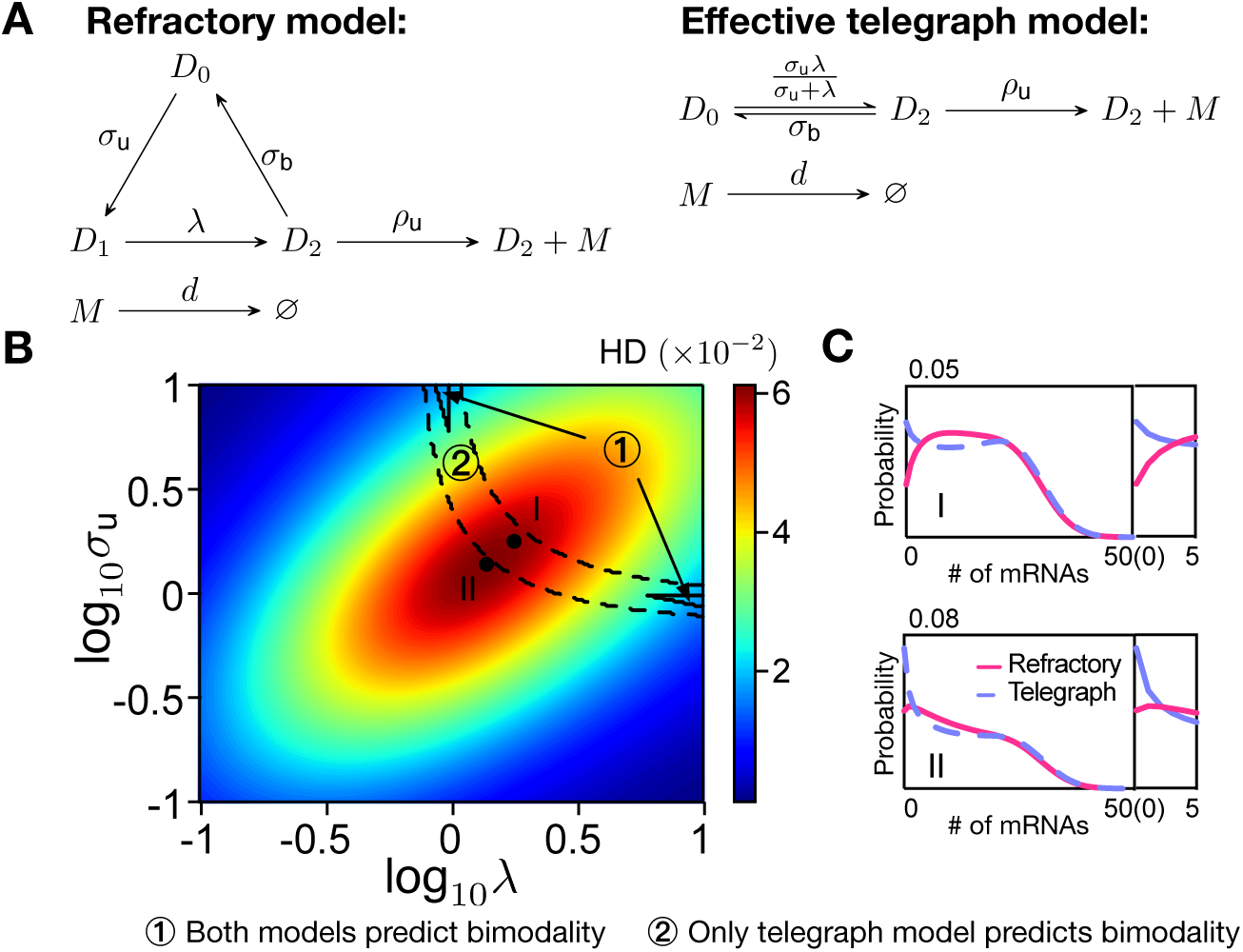
Effective telegraph model approximation for the refractory model. (A) Schematics of both models. (B) Hellinger distance between the steady-state distributions of mRNA numbers predicted by both models, and a bifurcation diagram of their number of modes(black lines) as a function of *σ*_u_ and *λ* with *σ*_b_ = 0.8, *ρ*_u_ = 30 and *d* = 1. (C) Distributions for Points I and II in (B), showing significant disagreement in the height of the zero mode (insets show a zoom at the mode at zero).

Given the reaction scheme illustrated in Fig. 4A, it follows that the temporal evolution of probability *P*_*θ*_(*n*) of finding *n* mRNAs and gene state *D*_*θ*_ (*θ* = 0, 1 or 2) can be described by the following master equations

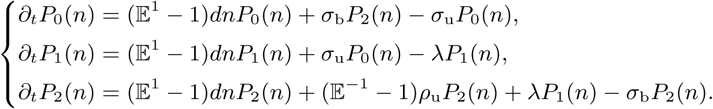

The corresponding generating function equations are given by

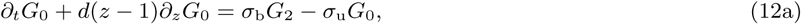

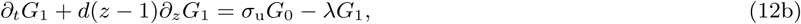

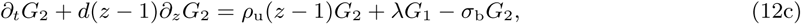

where *G*_*θ*_ = Σ_*n*_ *z*^*n*^ *P*_*θ*_(*n*). We intend to solve Eqs. (12) at steady state and thus set *∂*_*t*_*G*_*θ*_ = 0. Then, we solve *G*_1_ as a function of *G*_2_ from Eq. (12c), subsequently substitute it into Eq. (12b) and solve *G*_0_ as a function of *G*_2_. Following that, Eq. (12a) becomes an ordinary differential equation with *G*_2_ being the only variable to be solved

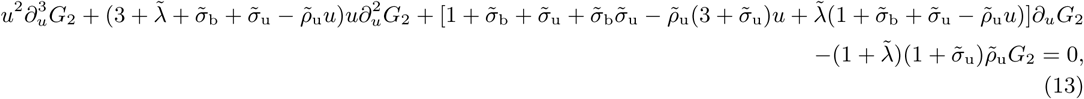

where 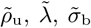 and 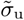 are the kinetic parameters normalized with respect to *d* and *u* = *z* − 1. Eq. (13) is the canonical form of the differential equation for the generalized hypergeometric function _2_*F*_2_, admitting the solution

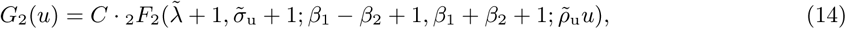

where *C* is an integration constant, and *β*_1_ and *β*_2_ denote

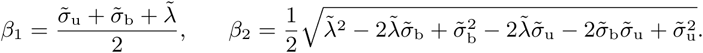

Summing Eqs. (12) leads to 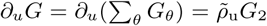, one can obtain *G* from Eq. (14) in the form of the generalized hypergeometric function:

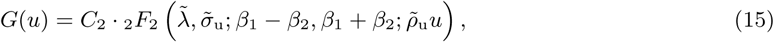

and *C*_2_ is found to be 1 by the normalization condition *G*(0) = 1. Eq. (15) together with 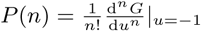 defines the distribution of mRNA numbers for the refractory model in steady-state conditions. A similar solution is also known for a generalization of the refractory model [29].

The next step is to map the refractory model onto an effective telegraph model by matching the mean mRNA numbers

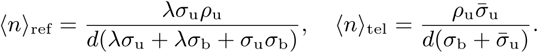

leading to an effective burst initiation rate 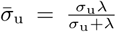 and the corresponding effective model shown in Fig. 4A right. Note that while the multi-scale model is approximately equivalent to an effective telegraph model with a renormalized mRNA production rate, the refractory model’s telegraph approximation leads to a renormalized rate of switching to the active state.

We then compare the steady-state distributions of the refractory model and its effective telegraph model. A heatmap of HD quantifying their distributional difference and a modality diagram (marked as black lines) of the two distributions are illustrated in Fig. 4B. Both the regions of high HD and Region 2 where only the telegraph model predicts bimodality are significantly large; also Region 1 where both predict bimodality is small. This shows that the refractory model, in general, is not well approximated by the telegraph model, particularly the latter’s probability for low mRNA numbers is not accurate – see Fig. 4C. Given the telegraph model’s excellent approximation to the multi-scale model, it is clear that the multi-scale model and refractory model can be distinguished.

### 2.6 Protein dynamics

Finally, for completeness, we extend the multi-scale model to provide analytic steady-state distributions of protein numbers. This allows interpretations of single-cell data of protein expression (see for example [30]). We consider the network in Fig. 1A with two additional reactions: (i) a first-order reaction modelling the translation of mRNA to proteins with rate constant *k*. (ii) a first-order reaction modelling the decay of protein with rate constant *d*_p_. It is shown in Appendix C that under the classic short-lived mRNA assumption (*d* ≫ *d*_p_) [20], the generating function corresponding to the steady-state distribution of protein numbers is given by

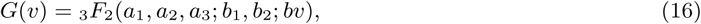

with *b*_1_ = (*σ*_b_ + *σ*_u_)*/d*_p_, *b*_2_ = (*σ*_b_ + *λ* + *ρ*)*/d*_p_, the mean translational burst size *b* = *k/d*, and the parameters *a*_1_, *a*_2_ and *a*_3_ being solutions of the equations

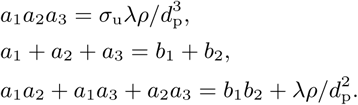

In the limit of large *λ* or *ρ*, we show in Appendix C that Eq. (16) reduces to the Gaussian hypergeometric function (_2_*F*_1_), which was reported in Ref. [20] for the classical three-stage model of gene expression in the limit of fast mRNA decay.

## 3 Conclusion

Here we performed the first detailed analytical study of a multi-scale model of bursty gene expression based on recent experimental data from mammalian cells [15]. The conventional telegraph model does not include an independently regulated pause release step and hence cannot differentiate the effects of changing polymerase pause release versus polymerase recruitment rates. Whereas, the multi-scale model studied here can distinguish these effects. While our model has three effective gene states (one of which regulating pause release), it is not a special case of existing multistate models because in our model, the gene state changes upon production of new nascent mRNAs to model the experimental observation that, unless the polymerase is unpaused (and nascent mRNA starts being actively transcribed by this polymerase), there can be no binding of new Pol II. In contrast, current models assume the gene state does not change upon production of mRNA because they model the production of a mature transcript without detailed modelling of the steps between transcriptional initiation and termination.

We have derived simple closed-form expressions for the approximate time evolution of the mRNA numbers and used the theory to understand which reactions contribute mostly to fluctuations. We also showed that (i) this model can be distinguished from the refractory model, another three gene state model popular in the literature. (ii) a number of previous models in the literature are special cases of our model, valid only in certain parameter regimes. Specifically the mRNA and protein distributions of the conventional three-stage model of gene expression provide a good approximation to the multi-scale bursting model in certain regions of parameter space as shown in Appendices B and C.

The simplicity of the equations for the mean and the variance allow the inference of rate parameters from single-cell data using maximum likelihood methods [31]. Potential extensions include (i) the impact of cell cycle effects such as binomial partitioning and variability in the cell cycle duration; (ii) introducing a detailed description of polymerase movement along the gene during elongation. The use of the recently developed linear mapping approximation [32] appears to be a promising means to extend the analytical solution of the present model to include feedback loops via DNA-protein interactions [33, 34].

An important result of the paper is that the time dependent mRNA distribution of the multi-scale model with polymerase dynamics and three states can be accurately approximated by the two state telegraph model, modified with a Michaelis-Menten like dependence of the effective transcription rate on polymerase abundance. Specifically, by Eq. (11) the transcription rate of a gene locus is *ρ*_u_ = *λρ/*(*λ* + *ρ*) where *λ* is the binding rate of Pol II (see Fig. 1A) which is proportional to the local number of Pol II molecules at the gene locus with active transcription [35]. This equation implies that the transcription rate is proportional to the local number of Pol II molecules if *λ* is approximately less than *ρ*, i.e. if the Pol II binding rate is less than or equal to the rate at which Pol II is unpaused. In contrast, if unpausing is the rate limiting step (*ρ* ≪ *λ*) then the transcription rate is practically independent of the local Pol II number.

Now when the number of gene copies doubles during replication, the local number of Pol II molecules will correspondingly decrease due to increased sharing of Pol II. Hence if we are in the regime *λ* ; ≲ *ρ*, the transcription rate per gene copy decreases; thus the total transcription rate for a gene per cell post-replication will be consequently slower than twice the total transcription rate pre-replication. This implies that the mean number of RNA per cell is not significantly affected by replication; indeed this “dosage compensation” has been observed experimentally for some genes in mouse embryonic stem cells [36] though a different explanation than above was suggested. In one study [37] it was estimated that for 6 yeast genes (RPB2, RPB3, TAF5, TAF6, TAF12, KAP104), the formation of the pre-initiation complex at the promoter (*λ*) is approximately equal to the rate at which the RNA polymerase escapes the promoter (*ρ*); hence gene dosage compensation via polymerase sharing, as implied by our model, maybe common. In contrast if we are in the regime *ρ* ≪ *λ*, the transcription rate per gene copy before and after replication is the same, and hence the total transcription rate for a gene per cell post-replication will be twice the total transcription rate pre-replication. This is also what is predicted by the telegraph model with constant burst initiation and termination rates, and observed experimentally for a reporter gene expressed from a strong synthetic promoter [36]. Note that since the mean burst size is the mean number of RNAs transcribed when the gene is on, by our reasoning above it also follows that when *λ* ; ≲ *ρ*, the mean burst size is altered upon gene replication. The idea that the number of RNA polymerases is the limiting factor in transcription has been recently hypothesized [38], and has implications for the mitigation of burden imposed by gene circuits in synthetic biology [39]. Our model here goes one step further by deriving the explicit relationship between the transcription rate and the number of RNA polymerases. Generally our model supports the observation that there are differences in transcriptional activity between different stages of the cell cycle [40] that cannot be explained by the conventional telegraph model.

## Author Contributions

Z.C. formulated the research question, performed the calculations, produced the figures and wrote an initial draft of the manuscript. T.F. performed some of the calculations for protein distributions. D.O. supervised the research and edited the manuscript. R.G. formulated the research question, supervised the research and wrote the manuscript with assistance from the co-authors.

## Acknowledgements

Z.C. gratefully acknowledges support of the UK Research Councils’ Synthetic Biology for Growth programme and of the BBSRC, EPSRC and MRC (B/M018040/1) and careful proofreading by J. Holehouse. R.G. acknowledges support from BBSRC grant no. BB/M025551/1. D.O. acknowledges support from the Human Frontier Science Program (grant no. RGY076/2015).

## Appendix

### A Analytic distribution for mRNA numbers when *ρ, λ* and *σ*_b_ are large

Given the large values of *ρ, λ* and *σ*_b_, we implement the following parametrization

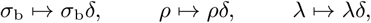

where *δ* is a large real number.

By means of the method of characteristics, solving Eq. (2) is tantamount to seeking a solution to the ODE system

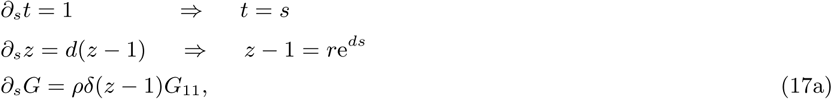

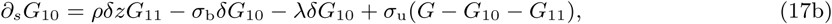

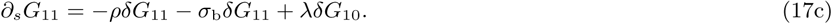

Dividing *δ* on both sides of Eqs. (17a)–(17c), one obtains a singular system consisting of

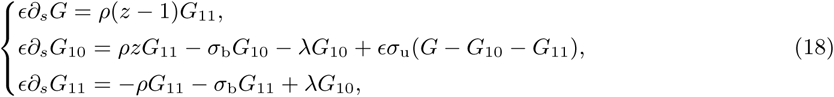

with *ϵ* = 1*/δ* ≃ 0. Expanding *G, G*_10_ and *G*_11_ in Eq. (18) as a series in powers of *ϵ*,

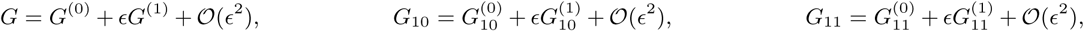

and matching the orders of *ϵ*, we have

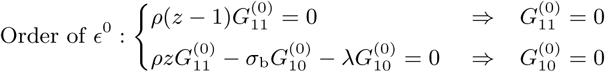

and

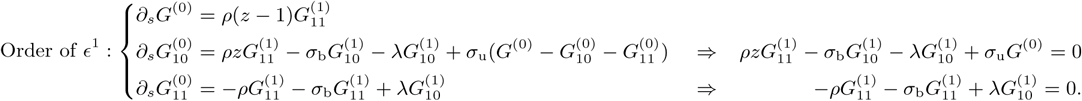

Then, we have

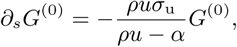

where *α* = *σ*_b_*γ*_2_*/λ* and *u* = *z* − 1 = *r*e^*ds*^. Its solution immediately follows as

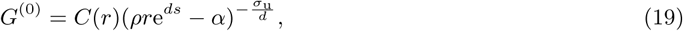

with *C*(*r*) being a function of *r* to be determined from the initial condition. Suppose that the initial condition for this process is *g*(*u*) = *G*^(0)^|_*t*=0_, which is known *a priori*. For instance, say the initial distribution of *n* mRNA molecules is *P* (*n*) = *p*_*n*_, then *g*(*u*) = Σ_*n*_ *p*_*n*_(*u* + 1)^*n*^. Letting *s* be equal to 0 (or equivalently *t* = 0), it follows *u* = *r* and *g*(*u*) = *g*(*r*), and we can establish the following relation

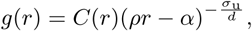

from which we can solve *C*(*r*) as

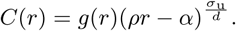

Substituting the latter back into Eq. (19) and replacing *r* = *u*e^−*dt*^, we can calculate the leading-order solution of *G* from (19) as

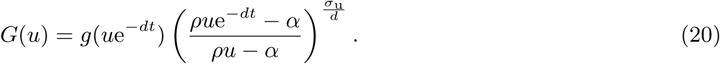

At steady state, the leading-order solution in (20) becomes

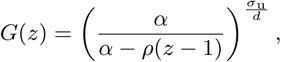

and the corresponding distribution of mRNA numbers is a negative binomial distribution NB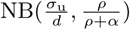.

### B Convergence to telegraph model for large *ρ*

To this end, we parametrize *ρ* as *ρ* ↦ *ρδ*, where *δ* is a large real number. As such, Eq. (2) can be recast as

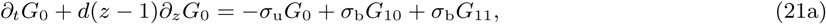

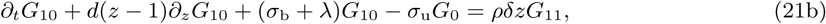

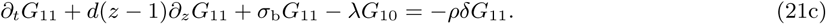

Dividing both sides of Eqs. (21b)-(21c) by *δ* and setting *ϵ* = *δ*^−1^, we have that

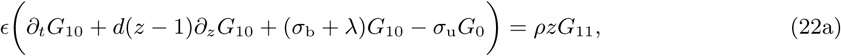

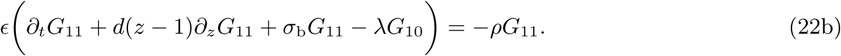

Again using the same method as before, we expand *G*_0_, *G*_10_ and *G*_11_ in Eqs. (21a) and (22) as a series in powers of, collect the terms for *ϵ*^0^ and *ϵ*^1^ and obtain

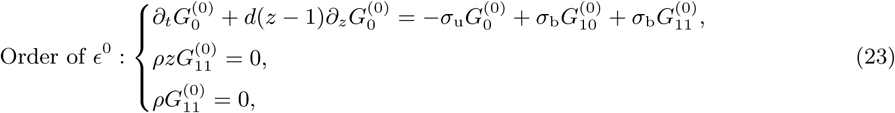

and

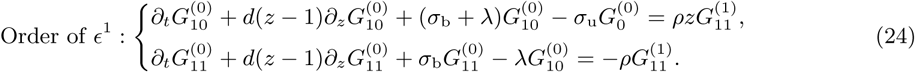

From Eq. (23), we can solve that 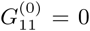, with which we can further get 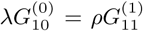 from Eq. (24). Given both results, Eqs. (23) and (24) can be simplified to

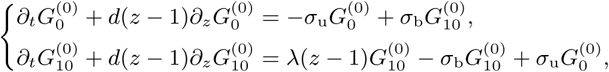

which are exactly the generating function equations of the telegraph model (See Eqs. (A2) and (A3) in [28]), thus showing that the multi-scale transcriptional bursting model converges to the telegraph model when *ρ* → ∞. A similar proof can be constructed to show that the telegraph model is also obtained in the limit *λ* → ∞.

### C Analytic marginal distribution for protein numbers for the multi-scale model in the limit of fast mRNA decay

To the reaction scheme illustrated in Fig. 1A, we add two reactions: (i) a first-order reaction modelling the translation of mRNA to proteins with rate constant *k*. (ii) a first-order reaction modelling the decay of protein with rate constant *d*_p_. The following coupled master equations describe the time evolution of the probability *P*_*θ*_(*n, m*) of finding *n* mRNAs, *m* proteins and gene state *D*_*θ*_ (*θ* = 0, 10, 11) in a cell:

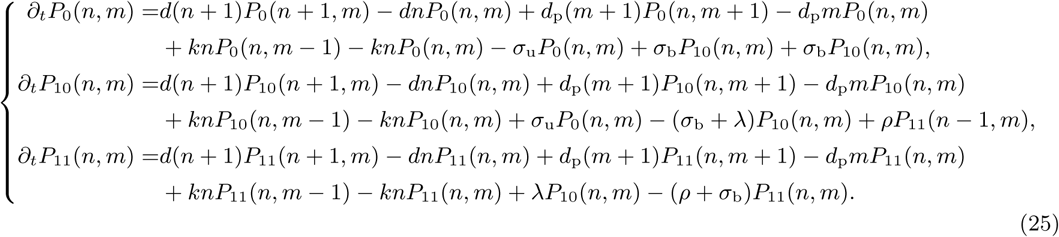

By defining *G*_*θ*_ = Σ_*n*_ Σ_*m*_ *z*_m_^*n*^ *z*_p_^*m*^ *P*_*θ*_(*n, m*), solving Eq. (25) is tantamount to seeking solutions to the set of differential equations

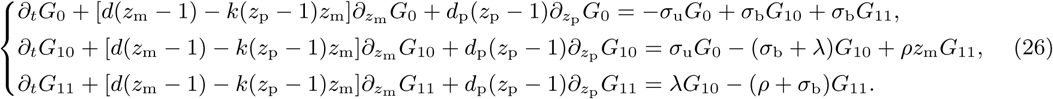

By means of the method of characteristics, Eq. (26) is equivalently represented as

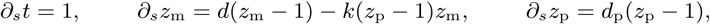

and

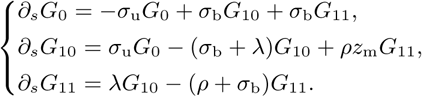

Assuming that mRNA decays much faster than protein such that *∂*_*s*_*z*_m_ ≃ 0 [20], we get that

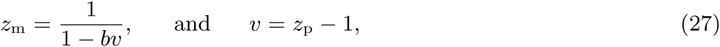

and *b* = *k/d* is the mean translational burst size. Using Eq. (27) we can reduce Eq. (26) to

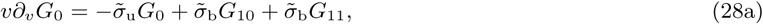

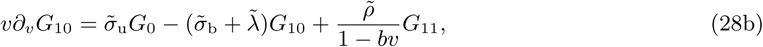

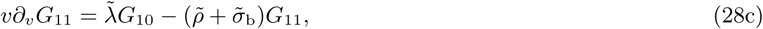

where 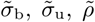 and 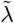 are kinetic parameters normalized with respect to protein degradation rate *d*_p_. It follows from summing Eqs. (28) that

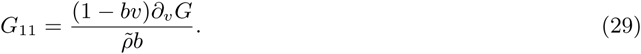

Using the definitions 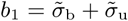 and 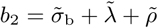 and plugging Eq. (29) into Eqs. (28b) and (28c), it gives us that

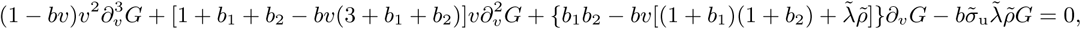

which admits a solution

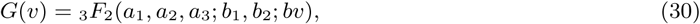

with *a*_1_, *a*_2_ and *a*_3_ being roots of

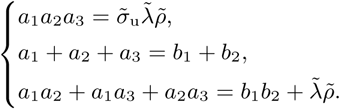

Hence summarizing, Eq. (30) and 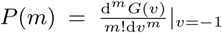 define the steady-state distribution of protein numbers, which is

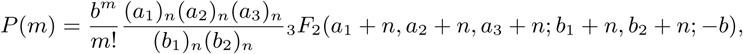

given that mRNA is short-lived.

Next we will show the solution Eq. (30) converges to the Gaussian hypergeometric function (_2_*F*_1_) for the three-stage gene expression model [20] when *ρ* is large. To this end, we parameterize 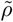 in Eqs. (28b)-(28c) as 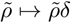 where *δ* is a large number. Dividing both sides of Eqs. (28b)-(28c) by *δ*, we have

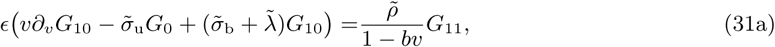

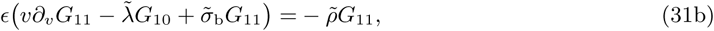

where *ϵ* = 1*/δ* ≃ 0. Again similarly, we expand *G*_0_, *G*_10_ and *G*_11_ in Eqs. (28a) and (31) as a series in powers of, collect the terms for *ϵ*^0^ and *ϵ*^1^ and obtain

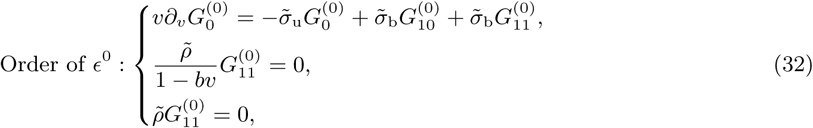

and

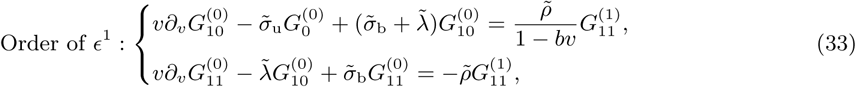

From Eqs. (32), we get 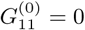, which is used to reduce Eqs. (33) and the first equation in Eq. (32) to

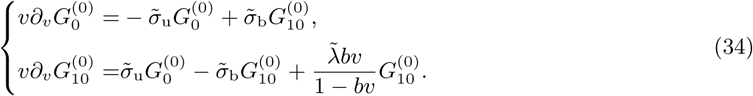

Note that Eq. (34), which is the leading order of Eqs. (28), is exactly the same as the generating functions of the three-stage gene expression model reported in [20] (See Eqs. (68)-(69) in SI thereof). By means of similar arguments, one can show the reduction of our model when *λ* is large.

